# Assessing the matrix effects of pigmented and non-pigmented salmon during multi-residue antibiotic analysis with liquid chromatography coupled to tandem mass spectrometry

**DOI:** 10.1101/2021.07.15.452301

**Authors:** Shiva Emami, Ameer Y. Taha

## Abstract

Several validated methods exist for the quantitation of antibiotics in seafood with ultra-high pressure liquid chromatography coupled with tandem spectrometry (UPLC-MS/MS). To our knowledge, none have explored the effects of co-eluting matrix components on the accuracy of quantitation. Such matrix effects could disproportionally change the ionization of analytes and their respective surrogate/internal standards during UPLC-MS/MS analysis, resulting in over-or under-estimation of antibiotic values. In this study, we measured matrix effects, alongside extraction recoveries for 30 antibiotics and their respective class-specific surrogate standards in Sockeye, King and Ivory (non-pigmented) salmon extracted using the QUEChERS method. A modified QUEChERS method involving dispersive or hydrophilic-lipophilic balance (HLB) solid phase extraction (SPE) was also tested on Sockeye salmon. Despite acceptable extraction recoveries for most antibiotics extracted using the QUEChERS method, significant matrix effects were observed for most antibiotic standards. Dispersive or HLB SPE clean-up did not improve analyte recoveries from Sockeye salmon, and in some cases, increased matrix effects. Accuracy and sensitivity were reduced when matrix effects were high. Our results demonstrate that matrix components in salmon cause matrix effects on antibiotics during UPLC-MS/MS analysis which could impact the accuracy and sensitivity of the analysis.

## Introduction

The use of antibiotics in aquaculture for therapeutic and/ or prophylactic purposes ^1^ has resulted in antibiotic contamination in seafood ^2,3^. Antibiotic residues in seafood pose a public health problem due to the potential for development of antibiotic resistance genes which could transfer to humans ^4,5^. In the US, antibiotic resistance is responsible for 35,000 premature deaths per year ^6^, which is why these residues are routinely measured in seafood by regulatory agencies.

The analysis of antibiotic residues in seafood is currently performed using ultra high-pressure liquid chromatography coupled to tandem mass spectrometry (UPLC-MS/MS) ^2,7-9^. Using tandem MS in multiple reaction monitoring (MRM) mode filters unwanted masses and enables simultaneous analysis of multiple antibiotic residues with high selectivity. A limitation of this approach, however, is that co-eluting components from the matrix could self-ionize or affect the ionization of the analytes, thus causing suppression or enhancement of the MS response. This phenomenon, known as the ‘matrix effect’ ^10^, may impact the sensitivity, accuracy and reproducibility of analysis, particularly when ion suppression or enhancement are not uniform across both the analyte and the surrogate (i.e. internal) standard used to quantify the analyte ^10,11^.

Salmon is one of the most popular seafoods consumed in U.S. and constitutes 14% of the total seafood consumption ^12^. The majority of the salmon consumed in the U.S. is farm-raised and antibiotic residues have been detected in farmed salmon samples collected from U.S. retail stores ^13^. In salmon, antibiotics are commonly extracted using non-polar solvents such as acetonitrile, with or without acid modifiers ^2,14,15^. Other methods include the QUEChERS (Quick, Easy, Cheap, Effective, Rugged and Safe) method ^7,9^ which was originally developed for pesticides ^16^ and later extended to antibiotic quantitation in various food matrices including seafood ^7-9^. QUEChERS also involves the use of acetonitrile in addition to a mixture of salts in water to drive the partitioning of antibiotics into the acetonitrile phase ^17^. QUEChERS followed by dispersive solid phase extraction (dispersive SPE) using sodium sulfate, primary, secondary amines (PSA), and C18 sorbents at a 900:50:150 ratio has been shown to minimize the matrix effects from seafood matrices ^7^. SPE involving hydrophilic-lipophilic balance (HLB) columns have also been used to extract antibiotics from fish 2. While, these methods have often shown good extraction recoveries for several antibiotic classes such as sulfonamides, quinolones and trimethoprim from salmon ^15^, there is no information about potential matrix effects from salmon matrix on antibiotics.

Salmon is particularly challenging compared to other types of fish because of its high carotenoid content. In other food matrices (e.g. fruit and vegetables), carotenoids are known to co-extract with other analytes of interest (e.g. pesticides) when using the QUEChERS method ^18^. One study showed that removing carotenoids from banana extracts using graphitized carbon black reduced matrix effects ^19^. Carotenoids such as astaxanthin are present in high amounts in salmonid muscle (3 to 38 mg/ Kg) ^20^, and they can potentially cause ion suppression or enhancement of co-extracted with antibiotics. Previous studies have shown that approximately 0.5% of salmon matrix components could be extracted into the final extract during QUEChERS extraction ^21^. Although small, these matrix components could cause significant ion suppression or enhancement as reported for other seafood matrices such as clam (*C. gallina*), mussel (*M. galloprovincialis*) and fish (*P. flesus*) when extracted using the QUEChERS method and analyzed by UPLC-MS/MS ^9^.

In the present study, we assessed the matrix effects of salmon, on antibiotics from eight classes commonly used in aquaculture following extraction by QUEChERS or QUEChERS plus dispersive SPE or HLB SPE. We hypothesized that will observe significant matrix effects of salmon with QUEChERS, and that based on other studies involving other fish matrices, dispersive or HLB SPE will improve sensitivity and reproducibility by minimizing matrix effects ^2,7^.

## Materials and methods

### Materials

Sockeye salmon (Open Nature Salmon Sockeye Alaskan Fillet, Wild caught) was purchased from a local supermarket in Davis, CA (USA). King salmon and Ivory King salmon were purchased from Savory Alaska (Leander, TX). LC/MS grade methanol and acetonitrile were obtained from Fisher Scientific (Hampton, NH, USA). Formic acid, sodium sulfate (NA_2_SO_4_) and sodium chloride (NaCl) were purchased from Sigma-Aldrich (St. Louis, MO). Trisodium citrate dehydrate (Alfa Aesar) was purchased from Fisher Scientific (Pittsburg, PA). Primary secondary amines (PSA) and C18 endcapped SPE bulk sorbents were purchased from Agilent technologies (Santa Clara, CA). SPE columns (1g, 20 cc cartridge, OASIS HLB) were purchased from Waters Corp. (Milford, MA). Antibiotic standards used in this study were from the following classes: **Amphenicols:** chloramphenicol (CAP), thiamphenicol (TAP), florfenicol (FF), florfenicol amine (FFA); **Tetracyclines:** tetracycline (TC), oxytetracycline (OTC), chlortetracycline (CTC), doxycycline HCl (DOX); **Sulfonamides:** sulfadimethoxine (SDM), sulfasalazine (SSZ), sufamethoxazole (SMX), sulfadiazine (SDZ); **Quinolones:** enrofloxacin (ENRO), flumequine (FLU), norfloxacin (NOR) and enoxacin (ENO); **Macrolides:** erythromycin (ERYTH), azithromycin (AZ), tylosin A (Tylosin), virginiamycin complex (VIRG-M1 and VIRG-S1), roxithromycin (ROX), tilmicosin phosphate (TILM); **B-lactams:** ampicillin anhydrous (AMP), penicillin G potassium salt (PEN-G), penicillin V (PEN-V) and amoxicillin (AMOX); **Lincosamides:** lincomycin (LIN), **Others:** trimethoprim (TRIM), ormetoprim (ORM) and.

AMOX (98%), ROX (97%), SDZ (99%), TAP (?) and FF (98%) were purchased from Fisher Scientific (Ward Hills, MA). ERYTH (94.8%) DOX HCl (98.8%), NOR (98%), AMP (99.6%), SDM (98.5%), ENRO (99.8%), TC (≥ 98%) and FFA (99.3%) were purchased from Sigma Aldrich (St. Louis, MO). CTC (98.0%), OTC (≥ 95%), FLU (100.0%), ENO (100%), AZ (99.5%), Tylosin (99.8%), VIRG (99.0%), PEN-G (99.5%), PEN-V (98.8%), SSZ (100%), SMX (100%), LIN (98%), TRIM (100%) and TILM (100%) were purchased from Cayman Chemicals (Ann Arbor, MI). CAP (98.5%) was purchased from Crescent Chemical (Islandia, NY). Isotopically labeled surrogates including CAP-D5 (chemical purity: 98%; isotopic purity: 98.3%), SMX-D4 (chemical purity: 98%; isotopic purity: 99.2%), sulfamethazine-D4 (SMZ-D4; chemical purity: 98%; isotopic purity: 95.9%), AZ-D3 (HPLC purity: 99.86%; isotopic purity: 93.9%), ERYTH-D6 (chemical purity: 95%; isotopic purity: 98.1%), TRIM-D3 (chemical purity: 99.49%; isotopic purity: 99.9%), LIN-D3 (chemical purity: 95%; isotopic purity: 99.6%), ENRO-D5 (HPLC purity: 99.61; isotopic purity: 99.40), ROX-D7 (HPLC purity: 96.04%; isotopic purity: 99.00%), L-(+)-AMP-D5 (chemical purity: 95%; isotopic purity: 99.00%), and ent-FFA-D3 (chemical purity: 98%; isotopic purity: 98.7%) were purchased from Toronto Research Chemicals (Toronto, Ontario, Canada). PEN-V-D5 (chemical purity: ≥98%; isotopic purity: ≥99%) was purchased from Cayman Chemicals (Ann Arbor, MI).

### Antibiotic stock solutions

Individual stock solutions of CAP, TAP, FF, FFA, TC, OTC, CTC, DOX, SDM, SMX, ENRO, ERYTH, AZ, VIRG, ROX, TILM, TRIM, ORM, LIN, CAP-D5, FFA-D3, SMX-D4, SMZ-D4, AZ-D3, ERYTH-D6, TRIM-D3, ROX-D7, ENRO-D5 were prepared in methanol at a concentration of 1 mg/mL. SSZ, SDZ, Tylosin, LIN-D3 were prepared at a concentration of 0.5 mg/mL in methanol and FLU, NOR and ENO were prepared at a concentration of 0.1 mg/mL in methanol to ensure their solubility. B-lactams along with their deuterated surrogates (APM-D5 and PEN-V-D5) were prepared in water (1 mg/mL).

Individual intermediate solutions of 10 µg/mL of each antibiotic were made in the same solvent as the stock solution and were used to prepare the antibiotics mixture solution (working mix). Working mixes were prepared in water: methanol 1:1 prior to the experiment.

### Sample preparation

Salmon samples were homogenized in dry ice using Sears solid state 10-speed blender at speed 7. The homogenates were stored in loose ziplock bags in a -20 °C freezer overnight (∼12 hours) to allow the dry ice to sublime.

### Antibiotic extraction using QUEChERS method

Antibiotics from salmon matrix were extracted using the QUEChERS method ^7^. Approximately, 1 g of wild caught Sockeye salmon homogenate was weighted and placed in 50 mL Falcon tubes (Fisher Scientific, cat # LS4541). In order to assess the recovery, samples were spiked with antibiotics working mix at a final concentrations of 20 ng/g per sample, by adding 100 µL of 200 ng/ml antibiotics working mix dissolved in water: methanol (1:1) (n=5). Control samples consisted of a non-spiked salmon matrix (n=1) and a method blank (n=1) spiked with deuterated surrogate standards mix (20 ng/g). The method blank did not contain sample but was extracted in a similar manner as the extraction for samples in the same tubes. To each tube, 8 mL Millipore water was added with five ceramic beads, to facilitate homogenization. Then, 30 mL of acetonitrile was added and samples were hand-shaken for about 10 seconds. 30 µL formic acid was added to each sample and they were shaken for 30 min at 200 rpm using the incubator shaker (New Brunswick Scientific, Excella E24 Incubator Shaker series). A salt mixture consisting of 4g Na_2_SO_4_, 1g NaCl and 1.5 g of trisodium citrate dihydrate was added to the samples and they were hand-shaken for about 10 seconds followed by mechanical shaking for 30 min at 200 rpm. The samples were then centrifuged at 3000 rpm for 10 min at 10 °C (SORVALL RT 6000D, rotor H1000B) and the supernatant layer (∼ 30 mL of acetonitrile) was transferred to new sets of 50 mL falcon tubes. The supernatant extract was dried under nitrogen. The extracts were reconstituted in 1 mL of water: methanol (1:1).

To assess matrix effects, 900 µL of water: methanol (1:1) was added to a representative sample that was spiked with 100 µL of 200 ng/mL antibiotic mix (i.e. 20 ng/mL in each sample) dissolved in water: methanol (1:1) after extraction (n=1). A second sample consisted of a standard only aliquot containing 900 µL of water: methanol (1:1) spiked with 100 µL of 200 ng/mL antibiotic mix (20 ng/mL; n=1). Samples were vortexed for 3 min and sonicated (Branson 1210, Danbury, CT) for 3 min for complete resuspension of the extracts. Samples were transferred to 2 mL centrifuge tubes (Sealrite, USA Scientific, FL), centrifuged at 12000 rpm (13523 ×g) for 2 min (Eppendorf, 5424 R),transferred to filter-containing centrifuge tubes (Ultrafree-MC-VV; PVDF 0.1 µm; Milipore Sigma, MA) and centrifuged for 10 min at 12000 rpm (13523 ×g). The last step was repeated if any visible residues were seen in the tubes. The extracts were transferred to LC vials (Phenomenex, CA) prior to UPLC-MS/MS analysis. Samples were analyzed on the same day of extraction.

#### Experiment 1. Effect of clean-up methods on antibiotic extraction recovery and matrix effects

Two clean-up procedures i.e. dispersive-SPE and SPE were performed following antibiotic extraction from Sockeye salmon using the QUEChERS method. Matrix effects were also tested. For the QUEChERS method followed by dispersive SPE, 6 mL of the 30 mL supernatant of the QUEChERS extract was transferred to 15 mL Falcon tubes containing Na_2_SO_4_/ PSA/C18 (900/50/150 mg). The tubes were mechanically shaken for 30 min at 200 rpm using incubator shaker (New Brunswick Scientific, Excella E24 incubator shaker series) and centrifuged at 3000 rpm for 10 min at 10 °C (SORVALL RT 6000D, rotor H1000B). Then, 3 Ml of the supernatant layer was taken and dried under nitrogen. Extracts were reconstituted in 100 µL of water: methanol (1:1).

For the QUEChERS method followed by SPE clean-up, the supernatant (∼30 mL) was diluted by bringing the total volume to 200 mL using milliQ Water. The pH was adjusted to 2.5 using 800 µL formic acid. Samples were then loaded onto OASIS HLB SPE columns (Waters, 20 cc; 1g) pre-conditioned with methanol (20 mL), pure water (6 mL) and pH= 2.5 water (6 mL). The cartridges were washed with milliQ water (10 mL) and dried under vacuum for 5 min. Antibiotics were eluted using 12 mL of methanol. The eluent was evaporated under nitrogen to dryness and reconstituted in 1 mL of water: methanol (1:1). The extracts were vortexed, sonicated, centrifuged and filtered as described above and analyzed using UPLC-MS/MS.

For matrix effects, 1 sample was spiked again after extraction (n=1) with 90 µL of water: methanol (1:1) and with 100 µL of 200 ng/mL antibiotic working mix dissolved in water: methanol antibiotic mix. Samples were vortexed for 3 min followed by 3 min sonication and then centrifuged and filtered as described above. Extracts were transferred to LC vials and analyzed on the same day using UPLC-MS/MS.

#### Experiment 2. Effect of matrix pigments (carotenoids) on antibiotic extraction recovery and matrix effects

The results from Experiment 1 revealed relatively high matrix effects form Sockeye salmon for most of antibiotics when they were extracted using the QUEChERS method and the matrix effects were not improved by SPE or dispersive SPE clean-up. Given that the Sockeye salmon used for method development in Experiment 1 has high amounts of carotenoids (astaxanthin; ∼ 38 mg/ Kg) ^20^, we hypothesized that carotenoids might be responsible for the observed matrix effects on antibiotics. To test this hypothesis, wild caught King salmon (red/orange color; representing salmon matrix containing carotenoids) and Ivory King salmon (ivory white color; representing salmon matrix without carotenoids) were spiked with antibiotics and extracted using the QUEChERS method as described in Experiment 1 (n= 5 per fish). Extracts were analyzed using UPLC-MS/MS and recovery and matrix effects were compared. We chose King salmon over Sockeye salmon for this experiment because the ivory white counterpart was available for this type of salmon but not for Sockeye salmon, allowing us to compare the effect of matrix carotenoids on antibiotics extraction.

### Instrumentation

Antibiotic analysis was performed using an Agilent ultra-high pressure liquid chromatography coupled to a 6460 Agilent triple quad (UPLC-MS/MS). Chromatographic separation of the antibiotic mixture was performed on AQUITY BEH C18 column (100 × 2.1 mm, 1.8 µm), using 0.1% formic acid in water (mobile phase A) and 0.1% formic acid in acetonitrile (mobile phase B) running at a flow rate of 0.3 mL/min and column temperature of 30 °C. The mobile phase gradient condition was as follows: initial time: 10% B, 8 min: 20% B, 11 min 60% B, 13 min 100% B, 15 min 100% B, 17 min: 10% B and 20 min: 10% B.

MS/MS analysis was performed using Agilent jet stream electrospray ionization (ESI) operating in both positive and negative mode as shown in Table 1. The acquisition method was dynamic multiple reaction monitoring (dMRM) scan type. The MS source parameters were as follows: sheath gas (nitrogen) temperature of 375 °C, sheath gas flow of 11 L/min, drying gas (nitrogen) temperature of 250 °C, nozzle voltage of 0V, nebulizer gas pressure of 40 psi and capillary voltage of 3500 V. Collision induced dissociation was carried out using nitrogen in the collision cell. Specific MS/MS parameters including precursor ions, fragmentor voltages, and product ions along with their specific collision energies for each compound are shown in Table 1.

**Table 1.**
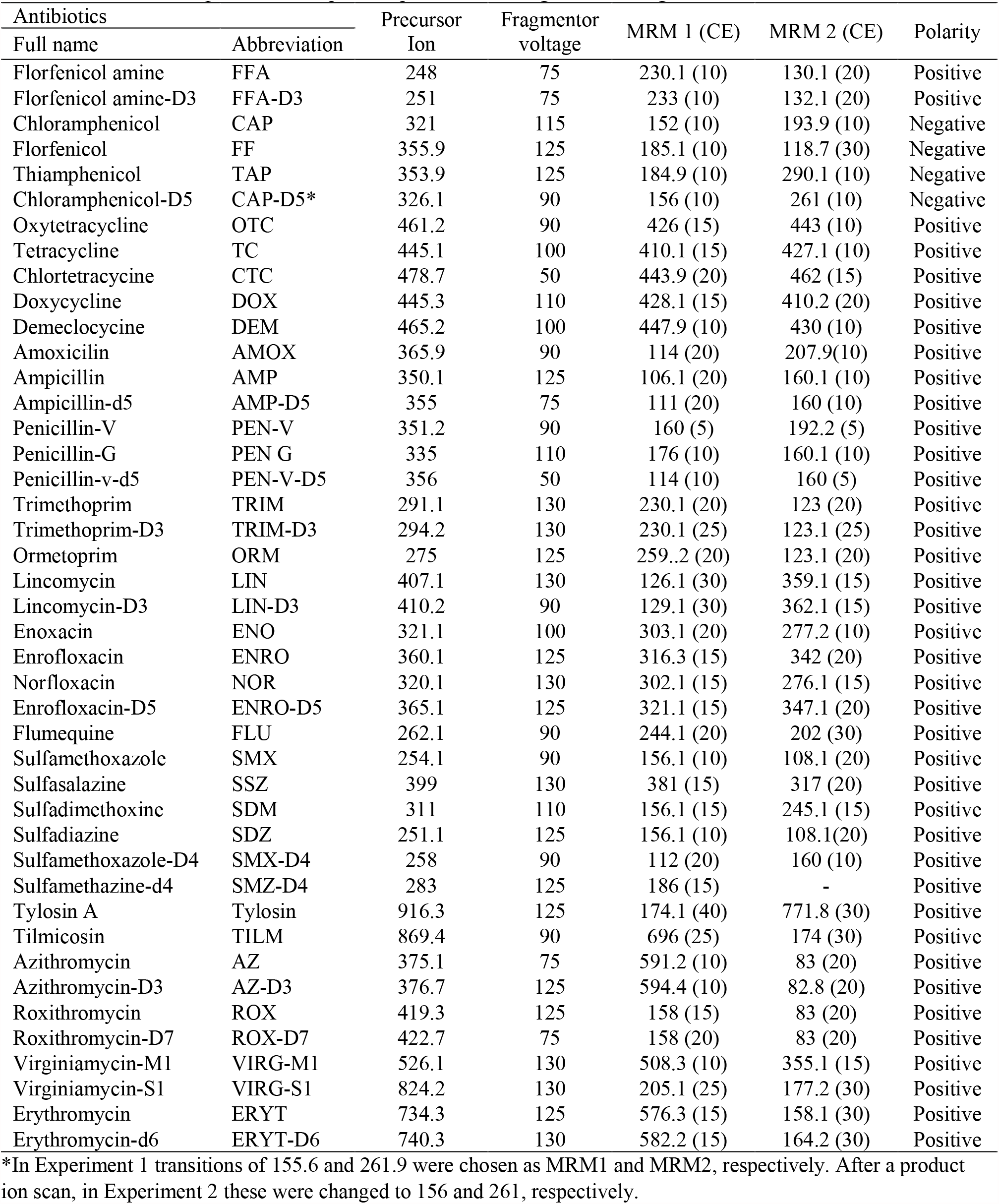
Precursor ion, quantifier and qualifier product ions, fragmentor voltage, and CE for antibiotic standards.

| Antibiotics |  | Precursor | Fragmentor | MRM 1 (CE) | MRM 2 (CE) | Polarity |
| --- | --- | --- | --- | --- | --- | --- |
| Full name | Abbreviation | Ion | voltage |  |  |  |
| Florfenicol amine | FFA | 248 | 75 | 230.1 (10) | 130.1 (20) | Positive |
| Florfenicol amine-D3 | FFA-D3 | 251 | 75 | 233 (10) | 132.1 (20) | Positive |
| Chloramphenicol | CAP | 321 | 115 | 152 (10) | 193.9 (10) | Negative |
| Florfenicol | FF | 355.9 | 125 | 185.1 (10) | 118.7 (30) | Negative |
| Thiamphenicol | TAP | 353.9 | 125 | 184.9 (10) | 290.1 (10) | Negative |
| Chloramphenicol-D5 | CAP-D5* | 326.1 | 90 | 156 (10) | 261 (10) | Negative |
| Oxytetracycline | OTC | 461.2 | 90 | 426 (15) | 443 (10) | Positive |
| Tetracycline | TC | 445.1 | 100 | 410.1 (15) | 427.1 (10) | Positive |
| Chlortetracycline | CTC | 478.7 | 50 | 443.9 (20) | 462 (15) | Positive |
| Doxycycline | DOX | 445.3 | 110 | 428.1 (15) | 410.2 (20) | Positive |
| Demeclocycline | DEM | 465.2 | 100 | 447.9 (10) | 430 (10) | Positive |
| Amoxicillin | AMOX | 365.9 | 90 | 114 (20) | 207.9(10) | Positive |
| Ampicillin | AMP | 350.1 | 125 | 106.1 (20) | 160.1 (10) | Positive |
| Ampicillin-d5 | AMP-D5 | 355 | 75 | 111 (20) | 160 (10) | Positive |
| Penicillin-V | PEN-V | 351.2 | 90 | 160 (5) | 192.2 (5) | Positive |
| Penicillin-G | PEN G | 335 | 110 | 176 (10) | 160.1 (10) | Positive |
| Penicillin-v-d5 | PEN-V-D5 | 356 | 50 | 114 (10) | 160 (5) | Positive |
| Trimethoprim | TRIM | 291.1 | 130 | 230.1 (20) | 123 (20) | Positive |
| Trimethoprim-D3 | TRIM-D3 | 294.2 | 130 | 230.1 (25) | 123.1 (25) | Positive |
| Ormetoprim | ORM | 275 | 125 | 259..2 (20) | 123.1 (20) | Positive |
| Lincomycin | LIN | 407.1 | 130 | 126.1 (30) | 359.1 (15) | Positive |
| Lincomycin-D3 | LIN-D3 | 410.2 | 90 | 129.1 (30) | 362.1 (15) | Positive |
| Enoxacin | ENO | 321.1 | 100 | 303.1 (20) | 277.2 (10) | Positive |
| Enrofloxacin | ENRO | 360.1 | 125 | 316.3 (15) | 342 (20) | Positive |
| Norfloxacin | NOR | 320.1 | 130 | 302.1 (15) | 276.1 (15) | Positive |
| Enrofloxacin-D5 | ENRO-D5 | 365.1 | 125 | 321.1 (15) | 347.1 (20) | Positive |
| Flumequine | FLU | 262.1 | 90 | 244.1 (20) | 202 (30) | Positive |
| Sulfamethoxazole | SMX | 254.1 | 90 | 156.1 (10) | 108.1 (20) | Positive |
| Sulfasalazine | SSZ | 399 | 130 | 381 (15) | 317 (20) | Positive |
| Sulfadimethoxine | SDM | 311 | 110 | 156.1 (15) | 245.1 (15) | Positive |
| Sulfadiazine | SDZ | 251.1 | 125 | 156.1 (10) | 108.1(20) | Positive |
| Sulfamethoxazole-D4 | SMX-D4 | 258 | 90 | 112 (20) | 160 (10) | Positive |
| Sulfamethazine-d4 | SMZ-D4 | 283 | 125 | 186 (15) | - | Positive |
| Tylosin A | Tylosin | 916.3 | 125 | 174.1 (40) | 771.8 (30) | Positive |
| Tilmicosin | TILM | 869.4 | 90 | 696 (25) | 174 (30) | Positive |
| Azithromycin | AZ | 375.1 | 75 | 591.2 (10) | 83 (20) | Positive |
| Azithromycin-D3 | AZ-D3 | 376.7 | 125 | 594.4 (10) | 82.8 (20) | Positive |
| Roxithromycin | ROX | 419.3 | 125 | 158 (15) | 83 (20) | Positive |
| Roxithromycin-D7 | ROX-D7 | 422.7 | 75 | 158 (20) | 83 (20) | Positive |
| Virginiamycin-M1 | VIRG-M1 | 526.1 | 130 | 508.3 (10) | 355.1 (15) | Positive |
| Virginiamycin-S1 | VIRG-S1 | 824.2 | 130 | 205.1 (25) | 177.2 (30) | Positive |
| Erythromycin | ERYT | 734.3 | 125 | 576.3 (15) | 158.1 (30) | Positive |
| Erythromycin-d6 | ERYT-D6 | 740.3 | 130 | 582.2 (15) | 164.2 (30) | Positive |

\*In Experiment 1 transitions of 155.6 and 261.9 were chosen as MRM1 and MRM2, respectively. After a product ion scan, in Experiment 2 these were changed to 156 and 261, respectively.

### Method characterization

Absolute recovery was calculated as follows:

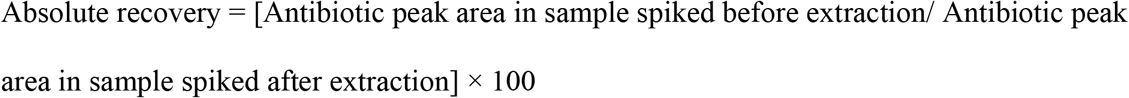

Matrix effects were calculated as follows:

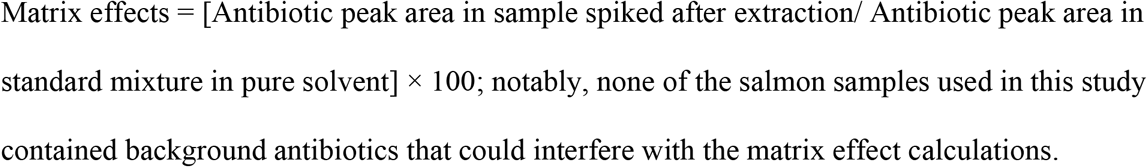

Accuracy was calculated according to equation below:

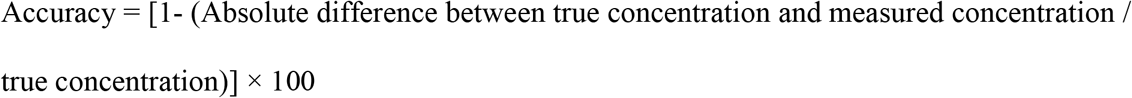

True concentration in the spiked samples was 20 ng/ mL for all antibiotics except for VIRG-M1 and VIRG-S1 which were spiked with 15 and 5 ng/ mL, respectively. This is because these two standards have been purchased as a single mixture at 75:25 ratio of VIRG-M1: VIRG-S1.

Antibiotic concentrations in spiked samples were calculated by the internal standard calibration method where surrogates were used to correct for both recoveries and matrix effects. A 9-point standard calibration curve (0.5-100 ng/ mL) containing a fixed amount of surrogate standard was made to derive the response factor. Calibration curves were generated by quadratic regression and 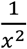 weighting factor was applied ^22^.

Method detection limits (MDL) were estimated following the procedure suggested by Environmental Protection Agency (EPA; 40 CFR, Appendix B to Part 136 revision 1.11, U.S.) by using the samples spiked with antibiotics:

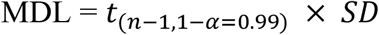

Where *t*_(*n*−1,1−α=0.99)_ is the student’s t value for 99% confidence level and degree of freedom of n-1, and SD represents standard deviation of the concentrations measured in the spiked salmon samples.

### Statistical analysis

Statistical analysis was performed using GRAPHPAD Prism 9.1.0 (La Jolla, CA, USA). In Experiment 1 (effect of clean-up methods on extraction recovery and matrix effects), analysis of variance (ANOVA) followed by Dunnett’s test was used to compare the results of each group with the control group i.e. QUEChERS extraction without clean-up. In Experiment 2 (effect of matrix pigments on extraction recovery and matrix effects), an unpaired t-test was used to compare the recoveries and matrix effects between the two fish matrices.

## Results

### Antibiotics choice and spike level

Thirty antibiotics from eight classes were selected for this study and are listed in **Table 1**. A representative MRM chromatogram of the 30 antibiotics spiked into the King salmon at 20 ng/g and extracted using the QUEChERS method alongside 13 deuterium labeled antibiotic surrogate standards is shown in **Figure 1**. The 30 antibiotics covered in this study constituted the majority of the antibiotics used in aquaculture farming in several countries ^23^, antibiotics banned for use in aquaculture in the U.S. (e.g. CAP) ^24^, and antibiotics that have been previously detected in seafood products in the U.S ^2^.

**Figure 1.**
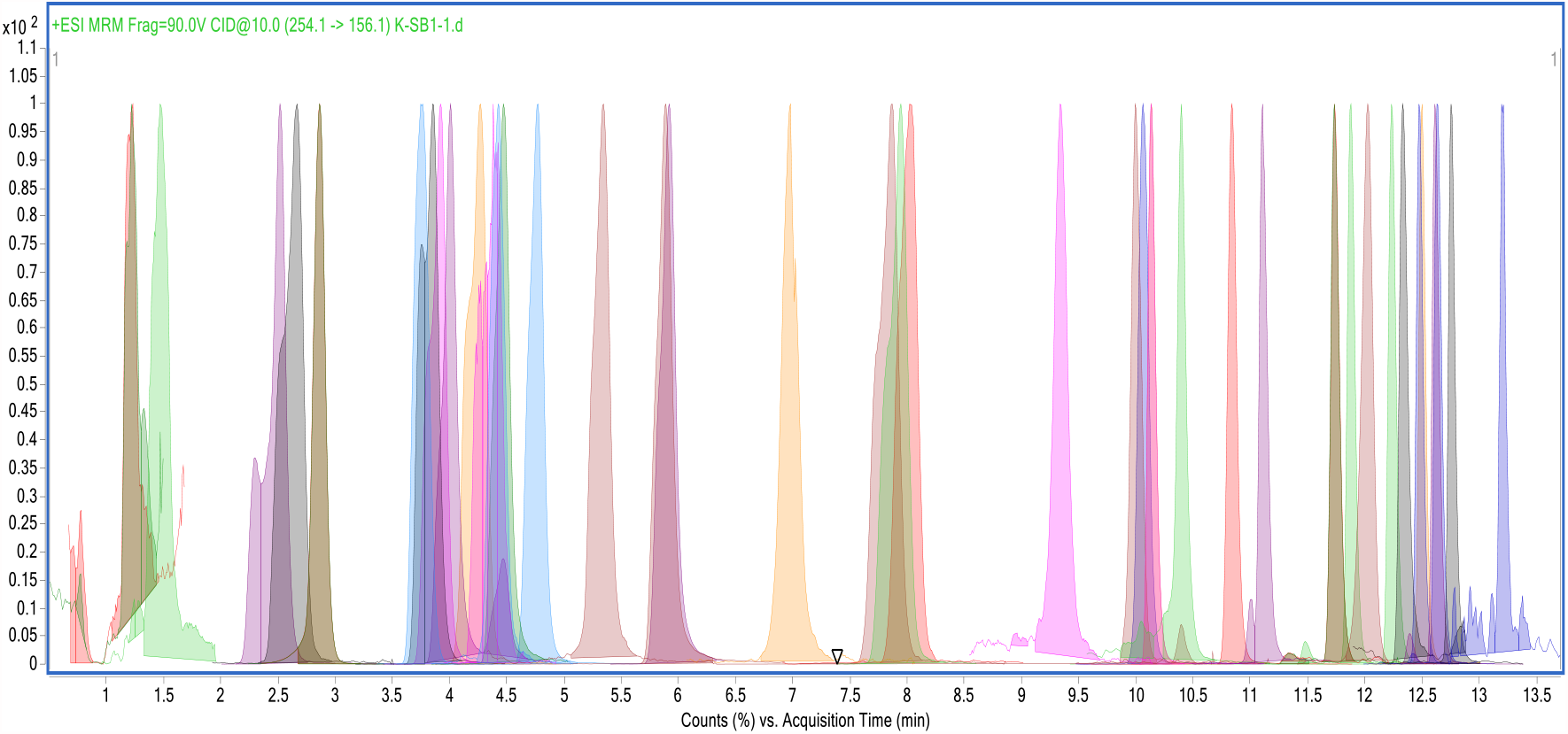
MRM chromatogram of target antibiotics spiked into King salmon matrix, extracted using QUEChERS (Experiment 2) and reconstituted in methanol: water (1:1)

#### Effect of clean-up methods on antibiotic extraction recovery and matrix effects

##### Antibiotic extraction recovery

The extraction recovery and matrix effects were tested in salmon spiked with ∼20 ng per g of sample. This level is less than half of maximum residue levels (MRLs) for most antibiotics (**Table S1**) ^25^.

**Table 2** shows the percent recoveries for antibiotics extracted using the QUEChERS method with or without subsequent dispersive or HLB SPE clean-up. The QUEChERS method resulted in mean percent recoveries of ∼56% for LIN and LIN-D3, ∼84% for TRIM, TRIM-D3 and ORM, 84-187% for quinolones, 70-96% for amphenicols, 42-61% for tetracyclines, 43-83% for sulfonamides, 19-97% for macrolides and 15-77% for B-lactams. AMOX from B-lactams class was not recovered at 20 ng/g spike level (i.e.it completely degraded during the extraction). Application of dispersive SPE following QUEChERS extraction did not significantly change the extraction recoveries for most of antibiotics (30 out of 42 unlabeled and labeled antibiotic standards), but significantly decreased the recovery of CAP-D5 (p < 0.05), FFA-D3 (p < 0.01), OTC (p < 0.05), AMP-D5 (p < 0.05), Tylosin (p < 0.05), VIRG-M1 (p < 0.0001), and significant increase for SMZ-D4 (p < 0.01), SDZ (p < 0.05), SMX-D4 (p < 0.01), SMX (p < 0.01), SDM (p < 0.01), ROX-D7 (p < 0.001), ROX (p < 0.001), and ERYTH-D6 (p < 0.0001) compared to QUEChERS (**Table 2**). When HLB SPE clean-up was applied after QUEChERS extraction, extraction recoveries significantly decreased for most antibiotics (36 out of 42 standards) compared to QUEChERS. FFA, SSZ and VIRG-S1 were not recovered from salmon matrix subjected to QUEChERS extraction followed by SPE clean-up at 20 ng/g spike level (**Table 2**).

**Table 2.** Percent recovery (mean ± SD; n=5) and percent matrix effects (n=1) of antibiotics extracted from Sockeye salmon spiked with 20 ng/g antibiotics using QUEChERS method or QUEChERS method followed by dispersive SPE or SPE using Oasis HLB columns. Ordinary one-way ANOVA followed by Dunnett’s post-hoc test was used to compare differences in recoveries of clean-up methods in comparison to QUEChERS only (as control group). When one group had non-detected values due to negligible recoveries, an unpaired t-test was used to compare the remaining two groups.

| Antibiotics class | Antibiotics | QUEChERS |  | QUEChERS-dSPE |  | QUEChERS-SPE |  |
| --- | --- | --- | --- | --- | --- | --- | --- |
|  |  | Recovery | Matrix effects | Recovery | Matrix effects | Recovery | Matrix effects |
| Lincosamides | LIN-D3 | 56 ± 6 | 34 | 54 ± 5 | 37 | 0.1 ± 0.1**** | 93 |
| | LIN | 57 ± 7 | 32 | 56 ± 5 | 34 | 0.2 ± 0.1**** | 90 |
| Trimethoprim and ormetoprim | TRIM-D3 | 84 ± 11 | 49 | 91 ± 9 | 50 | 4 ± 1**** | 105 |
| | TRIM | 83 ± 10 | 41 | 91 ± 9 | 43 | 4 ± 1**** | 90 |
| | ORM | 84 ± 10 | 47 | 92 ± 8 | 46 | 5 ± 2**** | 91 |
| Quinolones | ENRO-D5 | 142 ± 21 | 172 | 138 ± 14 | 181 | 16 ± 5**** | 357 |
| | ENO | 187 ± 42 | 187 | 155 ± 23 | 211 | 13 ± 5**** | 397 |
| | NOR | 180 ± 33 | 192 | 167 ± 18 | 166 | 17 ± 6**** | 432 |
| | ENRO | 143 ± 20 | 143 | 136 ± 12 | 150 | 16 ± 5**** | 295 |
| | FLU | 84 ± 9 | 15 | 91 ± 13 | 15 | 49 ± 4*** | 20 |
| Amphenicols | CAP-D5 | 96 ± 24 | 35 | 67 ± 15* | 46 | 51 ± 10** | 89 |
| | CAP | 92 ± 7 | 105 | 95 ± 6 | 102 | 63 ± 10*** | 137 |
| | FF | 93 ± 8 | 114 | 91 ± 6 | 117 | 54 ± 14**** | 163 |
| | TAP | 86 ± 10 | 118 | 93 ± 7 | 98 | 16 ± 6**** | 125 |
| | FFA-D3 | 74 ± 15 | 0.1 | 46 ± 10** | 0.3 | 1.3 ± 0.1**** | 38 |
| | FFA | 70 ± 23 | 0.1 | 57 ± 18 | 0.2 | ND | 39 |
| Tetracyclines | OTC | 47 ± 5 | 907 | 37 ± 6* | 576 | 3 ± 2**** | 1521 |
| | TC | 50 ± 6 | 860 | 42 ± 5 | 589 | 5 ± 3**** | 1289 |
| | CTC | 42 ± 4 | 526 | 32 ± 7 | 422 | 14 ± 8**** | 722 |
| | DOX | 61 ± 8 | 534 | 57 ± 6 | 370 | 31 ± 18** | 515 |
| Sulfonamides | SMZ-D4 | 47 ± 6 | 26 | 71 ± 16** | 36 | 10 ± 3*** | 115 |
| | SDZ | 55 ± 6 | 13 | 73 ± 13* | 17 | 11 ± 8**** | 109 |
| | SMX-D4 | 48 ± 4 | 38 | 69 ± 11** | 50 | 32 ± 5** | 132 |
| | SMX | 49 ± 4 | 26 | 69 ± 11** | 33 | 33 ± 5** | 87 |
| | SDM | 43 ± 5 | 11 | 68 ± 17** | 15 | 28 ± 3 | 23 |
| | SSZ | 83 ± 14 | 11 | 82 ± 9 | 14 | ND | 17 |
| B-Lactams | AMP-D5 | 17 ± 2 | 25 | 14 ± 2* | 36 | 0.3 ± 0.1**** | 117 |
| | AMP | 15 ± 2 | 42 | 12 ± 3 | 56 | 0.4 ± 0.2**** | 125 |
| | AMOX | ND | 1 | ND | 2 | 0.3 ± 0.1 | 121 |
| | PEN-V-D5 | 77 ± 19 | 5 | 75 ± 9 | 6 | 24 ± 5**** | 10 |
| | PEN-V | 75 ± 16 | 5 | 78 ± 11 | 6 | 23 ± 4**** | 10 |
| | PEN-G | 60 ± 17 | 11 | 77 ± 9 | 13 | 4 ± 1**** | 21 |
| Macrolides | AZ-D3 | 81 ± 11 | 212 | 84 ± 9 | 303 | 9 ± 2**** | 245 |
| | AZ | 79 ± 11 | 171 | 121 ± 61 | 232 | 9 ± 2* | 190 |
| | TILM | 82 ± 6 | 1285 | 86 ± 5 | 1006 | 50 ± 12**** | 871 |
| | ERYTH-D6 | 19 ± 18 | 23 | 87 ± 15**** | 31 | 2 ± 0.4 | 32 |
| | ERYTH | 19 ± 18 | 21 | 12 ± 2 | 202 | 2 ± 0.3* | 29 |
| | ROX-D7 | 87 ± 18 | 4 | 96 ± 14 | 3 | 44 ± 4*** | 6 |
| | ROX | 86 ± 17 | 3 | 93 ± 14 | 3 | 43 ± 5*** | 6 |
| | VIRG-M1 | 89 ± 3 | 8 | 66 ± 7**** | 7 | 58 ± 3**** | 8 |
| | VIRG-S1 | 97 ± 17 | 17 | 87 ± 20 | 11 | ND | 0 |
| | Tylosin | 76 ± 5 | 41 | 60 ± 7* | 42 | 47 ± 10*** | 47 |

ND: Not detected, \*p < 0.05, \*\* p < 0.01, \*\*\* p < 0.001, \*\*\*\* p < 0.0001 compared to QUEChERS.

##### Matrix effects

The QUEChERS method resulted in poor matrix effects in the form of ion suppression or enhancement for most antibiotics, where 100% indicates no matrix effects, < 70% indicates ion suppression and > 130% indicates ion enhancement. As shown in **Table 2**, ion suppression was observed after QUEChERS extraction for LIN and LIN-D3 (32-34%), TRIM, TRIM-D3 and ORM (41-49%), FLU (15%), CAP-D5 (35%), FFA and FFA-D3 (0.1%), sulfonamides (11-38%), B-lactams (1-42%) and macrolides (ERYTH, ERYTH-D6, ROX, ROX-D7, VIRG-M1, VIRG-S1 and Tylosin; 3-41%). On the other hand, quinolones including ENO, NOR, ENRO and ENRO-D5 (143-192%), tetracyclines (526-907%) and macrolides including AZ, AZ-D3 and TILM (171-1285%) showed ion enhancement. Matrix effects for amphenicols (CAP, FF and TAP) were within the optimal range (105-118%) following QUEChERS extraction.

QUEChERS followed by dispersive SPE did not improve matrix effects toward the optimal range of 70-130% for any of the ion suppressed or enhanced antibiotic standards. However, QUEChERS followed by SPE clean-up improved matrix effects for LIN and LIN-D3 (90-93%), TRIM, TRIM-D3 and ORM (91-105%), CAP-D5 (89%), Sulfonamides (SMZ-D4, SDZ, SMX, SMX-D4, 87-132%), AMP, AMOX and AMP-D5 (117-125%) (**Table 2**). On the other hand, ion enhancement was observed for CAP (137%) and FF (163%) following QUEChERS extraction and SPE clean-up.

#### Effects of matrix pigments (carotenoids) on antibiotic extraction recovery and matrix effects

##### Antibiotic extraction recovery

**Table 3** shows the percent recovery for antibiotics extracted by the QUEChERS method using two different matrices i.e. King salmon (with carotenoids) and Ivory King salmon (without carotenoids). Overall, the extraction recoveries were comparable between King salmon and Ivory King salmon. However, significant differences (p < 0.05) in the percent recovery of antibiotics were observed between the two types of fish for FFA-D3, LIN, LIN-D3, ENO, NOR, ENRO, ENRO-D5, SMX, SMX-D4, SDZ and SMZ-D4, which were higher by 6-14% in Ivory King salmon compared to King salmon. On the other hand, significantly higher recoveries (by 6-9%) were observed for AMP-D5, PEN-V and ROX in King salmon compared to Ivory King salmon (p < 0.05).

**Table 3.** Percent recovery (mean ± SD; n=5) and matrix effects (%, mean ± SD, n=5) of antibiotics extracted from King salmon (pink) and Ivory King salmon (white) spiked with 20 ng/g of antibiotics using the QUEChERS method. Unpaired t-test was used to compare differences in recoveries and matrix effects between two salmon matrices.

| Antibiotics class | Antibiotics | Recovery |  | Matrix effects |  |
| --- | --- | --- | --- | --- | --- |
|  |  | King Salmon | Ivory king Salmon | King Salmon | Ivory king Salmon |
| Lincosamides | LIN-D3 | 53 ± 3 * | 59 ± 4 | 69 ± 3 | 68 ± 6 |
| | LIN | 54 ± 3 * | 60 ± 4 | 70 ± 4 | 68 ± 6 |
| Trimethoprim and Ormetoprim | TRIM-D3 | 77 ± 5 | 83 ± 6 | 74 ± 2 | 77 ± 6 |
| | TRIM | 77 ± 4 | 83 ± 6 | 75 ± 2 | 76 ± 6 |
| | ORM | 82 ± 3 | 86 ± 6 | 93 ± 3 | 92 ± 8 |
| Quinolones | ENRO-D5 | 84 ± 5 * | 95 ± 5 | 639 ± 24 ** | 491 ± 65 |
| | ENRO | 84 ± 5 * | 94 ± 4 | 552 ± 26 ** | 451 ± 57 |
| | ENO | 71 ± 5 * | 78 ± 3 | 972 ± 31 *** | 771 ± 77 |
| | NOR | 74 ± 5 * | 83 ± 4 | 1083 ± 37 **** | 750 ± 72 |
| | FLU | 81 ± 5 | 77 ± 7 | 88 ± 6 * | 95 ± 3 |
| Amphenicols | CAP-D5 | 92 ± 8 | 97 ± 4 | 103 ± 2 | 102 ± 14 |
| | CAP | 102 ± 11 | 112 ± 13 | 159 ± 7 | 138 ± 24 |
| | FF | 110 ± 12 | 107 ± 10 | 118 ± 7 * | 100 ± 14 |
| | TAP | 133 ± 40 | 121 ± 22 | 83 ± 5 * | 71 ± 8 |
| | FFA-D3 | 41 ± 6 * | 53 ± 9 | 1 ± 0 | 1 ± 0 |
| | FFA | 44 ± 4 | 47 ± 10 | 1 ± 0 | 1 ± 0 |
| Tetracyclines | DEM | 42 ± 4 | 40 ± 3 | 525 ± 29 * | 606 ± 59 |
| | OTC | 54 ± 2 | 50 ± 5 | 328 ± 20 *** | 464 ± 47 |
| | TC | 49 ± 3 | 48 ± 4 | 354 ± 24 * | 404 ± 40 |
| | CTC | 44 ± 7 | 48 ± 4 | 396 ± 24 | 408 ± 31 |
| | DOX | 64 ± 4 | 59 ± 4 | 500 ± 39 *** | 679 ± 49 |
| Sulfonamides | SMZ-D4 | 59 ± 7 * | 71 ± 5 | 102 ± 4 | 90 ± 13 |
| | SMX-D4 | 57 ± 7 * | 69 ± 4 | 92 ± 2 | 86 ± 10 |
| | SMX | 58 ± 6 * | 70 ± 4 | 92 ± 1 | 87 ± 11 |
| | SDM | 53 ± 8 | 60 ± 7 | 131 ± 5 | 124 ± 12 |
| | SSZ | 81 ± 5 | 72 ± 7 | 49 ± 3 | 50 ± 2 |
| | SDZ | 48 ± 7 * | 62 ± 7 | 53 ± 3 | 53 ± 4 |
| B-Lactams | AMP-D5 | 24 ± 4 * | 18 ± 1 | 46 ± 1 | 46 ± 4 |
| | AMP | 27 ± 6 | 22 ± 3 | 52 ± 2 | 55 ± 4 |
| | AMOX | 14 ± 5 | 10 ± 2 | 15 ± 1 | 14 ± 2 |
| | PEN-V-D5 | 77 ± 3 | 70 ± 7 | 53 ± 3 | 56 ± 3 |
| | PEN-V | 79 ± 5 * | 70 ± 6 | 53 ± 4 | 56 ± 3 |
| | PEN-G | 46 ± 3 | 53 ± 8 | 67 ± 5 | 73 ± 5 |
| Macrolides | AZ-D3 | 79 ± 2 | 80 ± 7 | 333 ± 19 **** | 235 ± 16 |
| | AZ | 80 ± 2 | 80 ± 7 | 332 ± 20 **** | 236 ± 16 |
| | TILM | 88 ± 2 | 90 ± 6 | 530 ± 35 *** | 369 ± 53 |
| | ERYTH-D6 | 7 ± 2 | 9 ± 4 | 48 ± 15 * | 66 ± 8 |
| | ERYTH | 7 ± 2 | 9 ± 4 | 48 ± 15 * | 66 ± 8 |
| | ROX-D7 | 77 ± 4 | 71 ± 7 | 14 ± 1 * | 16 ± 1 |
| | ROX | 78 ± 4 * | 70 ± 7 | 13 ± 1 ** | 15 ± 1 |
| | VIRG-M1 | 66 ± 8 | 61 ± 10 | 47 ± 4 * | 53 ± 4 |
| | VIRG-S1 | 93 ± 22 | 70 ± 13 | 38 ± 4 | 43 ± 6 |
| | Tylosin | 72 ± 4 | 74 ± 7 | 57 ± 4 | 58 ± 4 |

\*p < 0.05, \*\* p < 0.01, \*\*\* p < 0.001, \*\*\*\* p < 0.0001; N/A: Not applicable.

##### Matrix effects

Significant differences in ion suppression and enhancement were observed between King and Ivory salmonfor 19 antibiotic standards, as shown in **Table 3**.

ERYTH (p < 0.05), ERYTH-D6 (p < 0.05), ROX (p < 0.01), ROX-D7 (p < 0.05) and VIRG-M1(p < 0.05) were less ion-suppressed (by 2-22%) in Ivory King salmon compared to King salmon; however, the matrix effect were below < 70% for these antibiotics in both salmon types. In Ivory King salmon, less ion suppression was observed for FLU (95% vs. 88%; p < 0.05), and more suppression for TAP (71 vs. 83%; p < 0.05) compared to King salmon; matrix effects were in optimal range (70-130%) for these antibiotics in both salmon types (**Table 3**).

Matrix effects in the form of ion enhancement were observed for FF (p < 0.05), quinolones (ENO (p < 0.001), NOR (p < 0.0001), ENRO (p < 0.01), ENRO-D5 (p < 0.01)) and macrolides (AZ (p < 0.0001), TILM (p < 0.001), AZ-D3 (p < 0.0001), which were significantly higher in Ivory King salmon compared to King salmon by 18-333% (**Table 3**). In addition, significantly more ion enhancement (50-179%) was observed in Ivory King salmon than King salmon for tetracyclines (OTC, TC (p < 0.05), DOX, DEM (p < 0.05). Matrix effects were out of the optimal range (70-130%) for all of these antibiotics except for FF for which matrix effects was in the optimal range (100 to 118%) in both salmon types.

##### Method accuracy

**Table 4** shows the accuracy of antibiotics quantified at the 20 ng/g spike level following extraction of Sockeye salmon using QUEChERS with or without subsequent clean-up, and spiked King salmon (containing carotenoid pigment)) and Ivory King salmon (non-pigmented) extracted with the QUEChERS method. Accuracies above 70% (% error < 30%) were considered acceptable. Using the QUEChERS method in Sockeye salmon, poor accuracies (<70%) were obtained for 14 antibiotics, including FLU, amphenicols (CAP, FF, TAP, FFA), AMP, PEN-G, VIRG-M1, VIRG-S1, TILM, Tylosin, SDM, SSZ and SDZ. Dispersive SPE or HLB SPE following the QUEChERS extraction did not improve accuracy, except for FFA, VIRG-M1 and Tylosin where dispersive SPE improved the method accuracy to 81-87%. Accuracy was also reduced for AZ extracted with QUEChERS and cleaned with dispersive SPE (69%) compared to QUEChERS without clean-up (83%). The accuracies were not calculated for tetracyclines in Sockeye salmon because of the lack of proper surrogate (unlabeled surrogate DEM was used) at the time of experiment.

**Table 4.** Accuracy (%) of target antibiotics in salmon matrices (Sockeye, King and Ivory King). For Sockeye salmon three extraction methods i.e. QUEChERS, QUEChERS-dSPE and QUEChERS-SPE were tested. King and Ivory King salmon were extracted using QUEChERS only. Data are represented as mean ± SD (n=5). Accuracy values below 70% were considered poor and marked with ǂ. Accuracies for OTC, TC, CTC and DOX were not calculated in Sockeye salmon matrix due to the lack of proper surrogate at the time of the experiment.

| Antibiotic | Tagged surrogate | Sockeye Salmon |  |  | King Salmon | Ivory King Salmon |
| --- | --- | --- | --- | --- | --- | --- |
|  |  | QUEChERS | QUEChERS-dSPE | QUEChERS-SPE | QUEChERS | QUEChERS |
| LIN | LIN-D3 | 99 ± 0 | 99 ± 1 | 96 ± 3 | 98 ± 0 | 98 ± 0 |
| TRIM | TRIM-D3 | 91 ± 1 | 92 ± 1 | 97 ± 1 | 96 ± 1 | 98 ± 0 |
| ORM | TRIM-D3 | 94 ± 1 | 97 ± 1 | 63 ± 4 † | 66 ± 4 † | 79 ± 3 |
| FLU | TRIM-D3 | 30 ± 1 † | 31 ± 3 † | -32 ± 59 † | 70 ± 14 | 86 ± 7 |
| ENO | ENRO-D5 | 88 ± 9 | 91 ± 4 | 72 ± 6 | 68 ± 5 † | 63 ± 3 † |
| NOR | ENRO-D5 | 86 ± 6 | 88 ± 3 | 96 ± 3 | 29 ± 8 † | 31 ± 2 † |
| ENRO | ENRO-D5 | 90 ± 1 | 86 ± 1 | 95 ± 1 | 85 ± 1 | 86 ± 1 |
| CAP | CAP-D5 | -165 ± 64 † | -121 ± 43 † | 2 ± 57 † | 37 ± 7 † | 39 ± 12 † |
| FF | CAP-D5 | -163 ± 57 † | -126 ± 44 † | -9 ± 74 † | 75 ± 7 | 81 ± 7 |
| TAP | CAP-D5 | -85 ± 74 † | -78 ± 36 † | 51 ± 26 † | 81 ± 15 | 91 ± 9 |
| FFA | FFA-D3 | 66 ± 17 | 81 ± 22 | ND | 88 ± 2 | 83 ± 13 |
| OTC | DEM | N/A | N/A | N/A | 86 ± 5 | 81 ± 6 |
| TC | DEM | N/A | N/A | N/A | 84 ± 6 | 84 ± 1 |
| CTC | DEM | N/A | N/A | N/A | 85 ± 6 | 89 ± 3 |
| DOX | DEM | N/A | N/A | N/A | 82 ± 6 | 85 ± 4 |
| AMP | AMP-D5 | 64 ± 8 † | 64 ± 14 † | 64 ± 4 † | 72 ± 5 | 56 ± 10 † |
| AMOX | AMP-D5 | ND | ND | 67 ± 33 † | 21 ± 4 † | 17 ± 3 † |
| PEN-V | PEN-V-D5 | 90 ± 8 | 96 ± 3 | 96 ± 4 | 98 ± 1 | 99 ± 1 |
| PEN-G | PEN-V-D5 | 35 ± 11 † | -13 ± 7 † | 46 ± 8 † | 75 ± 7 | 94 ± 5 |
| VIRG-M1 | PEN-V-D5 | 39 ± 32 † | 82 ± 6 | -20 ± 43 † | 72 ± 6 | 78 ± 5 |
| VIRG-S1 | PEN-V-D5 | 63 ± 39 † | 61 ± 7 † | -36 ± 81 † | 72 ± 13 | 64 ± 13 † |
| AZ | AZ-D3 | 83 ± 3 | 69 ± 46 † | 85 ± 4 | 95 ± 1 | 94 ± 1 |
| TILM | AZ-D3 | -300 ± 35 † | -152 ± 21 † | -1016 ± 77 † | 34 ± 3 † | 56 ± 8 † |
| ERYTH | ERYTH-D6 | 96 ± 3 | 98 ± 1 | 96 ± 1 | 98 ± 1 | 98 ± 2 |
| ROX | ROX-D7 | 93 ± 2 | 97 ± 1 | 96 ± 2 | 90 ± 1 | 91 ± 2 |
| Tylosin | SMX-D4 | 21 ± 8 † | 87 ± 3 | 61 ± 7 † | 80 ± 6 | 72 ± 6 |
| SMX | SMX-D4 | 76 ± 1 | 75 ± 2 | 75 ± 1 | 98 ± 2 | 98 ± 1 |
| SDM | SMX-D4 | 31 ± 2 † | 33 ± 3 † | 16 ± 1 † | 72 ± 8 | 74 ± 10 |
| SSZ | PEN-V-D5 | -27 ± 37 † | -36 ± 12 † | ND | 96 ± 3 | 95 ± 2 |
| SDZ | SMZ-D4 | 65 ± 3 † | 55 ± 2 † | 77 ± 18 | 45 ± 4 † | 53 ± 3 † |

ND: Not detected; N/A: Not applicable

**Table 5.** Method detection limits (MDLs, ng/g fish) of the target antibiotics in Sockeye, King and Ivory King salmon. In Sockeye salmon MDLs were calculated for the three extraction methods including QUEChERS, QUEChERS-dSPE and QUEChERS-SPE.

| Antibiotics | Sockeye Salmon |  |  | King<br>Salmon | Ivory King<br>Salmon |
| --- | --- | --- | --- | --- | --- |
|  | QUEChERS | QUEChERS-dSPE | QUEChERS-SPE | QUEChERS | QUEChERS |
| LIN | 0.56 | 0.63 | 2.02 | 0.35 | 0.34 |
| TRIM | 1.00 | 0.47 | 1.02 | 0.84 | 0.20 |
| ORM | 0.88 | 0.38 | 2.98 | 3.19 | 1.96 |
| FLU | 0.98 | 2.09 | 43.85 | 10.49 | 5.44 |
| ENO | 7.09 | 3.23 | 4.39 | 3.81 | 2.53 |
| NOR | 4.50 | 2.39 | 2.13 | 6.29 | 1.37 |
| ENRO | 1.04 | 0.95 | 0.86 | 0.55 | 0.42 |
| CAP | 48.24 | 32.20 | 42.90 | 5.28 | 8.89 |
| FF | 42.75 | 33.19 | 55.47 | 5.59 | 4.88 |
| TAP | 55.44 | 27.04 | 19.77 | 17.97 | 10.21 |
| FFA | 13.07 | 17.31 | N/A | 8.52 | 10.02 |
| OTC | N/A | N/A | N/A | 3.65 | 4.12 |
| TC | N/A | N/A | N/A | 4.25 | 0.98 |
| CTC | N/A | N/A | N/A | 4.77 | 2.37 |
| DOX | N/A | N/A | N/A | 4.76 | 2.77 |
| AMP | 5.72 | 10.82 | 3.22 | 4.08 | 7.84 |
| AMOX | N/A | N/A | 36.60 | 2.74 | 2.12 |
| PEN-V | 9.02 | 1.91 | 4.06 | 1.67 | 1.22 |
| PEN-G | 7.91 | 5.59 | 6.14 | 5.35 | 3.90 |
| VIRG-M1 | 18.02 | 3.62 | 23.95 | 3.27 | 3.05 |
| VIRG-S1 | 8.45 | 1.33 | 31.85 | 2.48 | 2.38 |
| AZ | 2.40 | 41.48 | 9.63 | 0.85 | 0.44 |
| TILM | 26.05 | 15.56 | 57.63 | 2.35 | 5.83 |
| ERYTH | 2.10 | 1.14 | 2.61 | 1.37 | 2.23 |
| ROX | 1.52 | 2.45 | 1.57 | 1.00 | 1.68 |
| Tylosin | 6.01 | 2.29 | 5.35 | 4.46 | 4.72 |
| SDZ | 2.30 | 1.33 | 23.55 | 2.89 | 2.35 |
| SMX | 0.50 | 1.19 | 0.40 | 1.57 | 0.76 |
| SDM | 1.29 | 2.24 | 0.86 | 6.32 | 7.72 |
| SSZ | 27.85 | 9.01 | N/A | 3.37 | 2.65 |

689 N/A: not applicable

The QUEChERS method enabled accurate (accuracy > 70%) quantitation of 24 out of 30 antibiotics at 20 ng/g fish spike level in King Salmon and 22 out of 30 antibiotics in Ivory King salmon. The antibiotics with poor quantitation accuracies included ENO, NOR, CAP, AMOX, TILM, and SDZ for both King and Ivory King salmon matrices. ORM in King salmon, and AMP and VIRG-S1 in Ivory King salmon also showed less than 70% accuracy in quantitation.

Overall, the data suggest that accuracy varies depending the type of salmon analyzed. Measurements were least accurate for Sockeye salmon, followed by Ivory King Salmon and King Salmon.

##### Method detection limit (MDL)

MDLs ranged from 0.56 ng/g for LIN to 48.24 ng/g for CAP in Sockeye salmon extracted using the QUEChERS method. Using clean-up methods after the QUEChERS did not affect the MDLs in a consistent manner. In some cases, SPE clean-up increased the MDLs.

In King and Ivory King salmon MDLs ranged from 0.35 ng/g for LIN to 10.21-17.97 ng/g for TAP. MDLs were generally lower in Ivory King salmon than King Salmon.

## Discussion

In this study, 30 antibiotics from eight classes were assessed for their extraction recoveries (from salmon) and matrix effects using the QUEChERS method with or without clean-up steps. Acceptable recoveries were observed when using the QUEChERS method to extract antibiotics from Sockeye salmon. However, matrix effects in the form of ion suppression or enhancement were observed for most analytes. Dispersive SPE clean-up following QUEChERS extraction did not eliminate the matrix effects for antibiotics in Sockeye salmon, and in many cases reduced extraction recoveries. SPE clean-up with HLB columns improved matrix effects for some antibiotics (LIN, LIN-D3, TRIM, TRIM-D3, ORM, SMZ-D4, SMX-SMX-D4, SDZ, AMP, AMP-D5), but reduced extraction recoveries for most antibiotics (**Table 2**). The absence of carotenoids from Ivory salmon slightly increased the extraction recoveries for 11 antibiotics and improved the matrix effects for 14 antibiotics compared to King salmon containing carotenoids, although improvements in matrix effects remained below the optimal 70-130% range. Conversely, greater matrix effects were obtained for tetracyclines and TAP when carotenoids were not present in matrix, i.e. in Ivory King salmon compared to King salmon (**Table 3**). Overall, our findings demonstrate varying degrees of antibiotic extraction recoveries and matrix effects in Sockeye, King and Ivory salmon that are unrelated to pigment (carotenoid) content.

The QUEChERS method resulted in acceptable extraction recoveries for most antibiotics at 20 ng/g spike level in Sockeye salmon. 21 out of 42 showed recoveries > 70% and 12 out of 42 analytes showed extraction recoveries of 40-69% (**Table 2**). Recoveries > 30% are usually considered acceptable in multi-residue extraction methods due to the challenges of dealing with antibiotic-matrix interactions ^9^. AMOX was not detected at 20 ng/g spike level and AMP and ERYTH were the only antibiotics showing extraction recoveries below 30% (7-27%). This could be due to their degradation during formic acid acidification at the beginning of the extraction. Degradation of some B-lactams (e.g. AMOX) in the presence of 0.2 or 1% formic acid in acetonitrile has been previously reported ^14^. Also, at low pH, ERYTH could transform to other metabolites such as anhydro-ERYTH and ERYTH-enol ether ^26^.

Optimal antibiotic spike recoveries were observed in Sockeye salmon extracts, except for quinolones including ENO, NOR, ENRO and ENRO-D5, which showed extraction recoveries above 100% (142-187%). This is likely due to adsorption of this class of antibiotics to the glass vial containing the working mix solution ^27^. Adsorption could reduce antibiotic levels in the samples spiked after extraction compared to samples spiked before extraction (thus yielding a calculated extraction recovery value above 100%).

The QUEChERS method resulted in significant matrix effects on antibiotics extracted from Sockeye salmon, despite optimally extracting most antibiotics with high recoveries from this matrix. The majority of compounds showed reduced MS response (i.e. ion suppression) in salmon compared to the pure solvent, except for tetracyclines, quinolones (ENO, NOR, ENRO, ENRO-D5) and macrolides (AZ, AZ-D3 and TILM) which showed enhanced the MS response, suggesting ion enhancement (**Table 2**). Enhanced instrument response for AZ, TILM and TC has been reported in other seafood matrices ^9^. In addition, antibiotics showing enhanced MS response such as quinolones and tetracyclines were reported to interact with the glass (vial) surface ^27,28^ and organic/ inorganic matter from food and environmental matrices ^29-31^. It is possible that antibiotics sorption onto the glass is reduced in the salmon extract due to the preferred affinity of some compounds to the dissolved matrix components, resulting in enhanced response in the salmon extract compared to the pure solvent.

Application of dispersive SPE clean-up using the Na_2_SO_4_/PSA/ C18 (900/50/150) sorbents did not improve matrix effects (**Table 2**). Na_2_SO_4_ is used to absorb trace amounts of water left in acetonitrile phase, and PSA and C18 are expected to remove polar lipids, such as fatty acids and phospholipids, and non-polar lipids from fish matrix, respectively ^32^. The inefficiency of the dispersive SPE in improving matrix effects suggests that other interferences from Sockeye salmon matrix might be the cause of ion suppression or enhancement.

The other clean-up method, i.e. SPE using OASIS HLB cartridges, improved matrix effects for some of the ion suppressed/ enhanced antibiotics, however this was accompanied by reduced extraction recoveries for all of the tested antibiotics (**Table 2**) suggesting that the SPE column retained both interfering matrices and antibiotics without separating them. Alternatively, antibiotic loss during SPE might be due to column overloading with other matrix components^33^ such as lipids and carotenoids, thus reducing the accessibility of active sites available for antibiotic binding. The retention of carotenoids in the column was evident from the orange/ red color of the column material during extraction. The ability of the column to retain carotenoids and lipids could explain the improved matrix effects observed here (i.e. whatever antibiotic escaped the column, did so alone and without interfering matrix components such as carotenoids).

The absence of carotenoids in Ivory King salmon, versus King salmon (with carotenoids), resulted in improved matrix effects for 14 out of 30 antibiotic standards. However, matrix effects for these compounds still remained out of optimal range (70-130%). In addition, reduced matrix effects were observed for tetracyclines, FF and TAP (**Table 3**). This suggests that carotenoids at the levels present in King salmon are not major contributors to the matrix effects. On the other hand, King salmon has lower content of carotenoids compared to Sockeye salmon (5.4 mg of astaxanthin per kg of flesh in King salmon vs. 28-36 mg/kg in Sockeye salmon)^20^. Therefore, it is still possible that carotenoids when found at high levels in matrices such as Sockeye salmon, might cause matrix effects during antibiotic analysis with UPLC-MS/MS.

Accuracy was impacted by the observed matrix effects. For instance, in Sockeye salmon, high matrix effects were observed for most antibiotic standards (**Table 2**) which is why only 11 out of 26 antibiotics were accurately quantifiable (accuracy > 70%). However, this value increased to 24 out of 30 antibiotics in King salmon and 22 out of 30 antibiotics in Ivory King salmon (**Table 4**), which both showed improved matrix effects compared to the Sockeye salmon (**Table 3**). More specifically, FLU, SDM, SSZ, PEN-G, VIRG-M1 and VIRG-S1 which showed high matrix effects following extraction from Sockeye salmon using QUEChERS method (8-17%), were not accurately quantified (accuracy < 70%) (**Table 2 and Table 4**). On the other hand, these antibiotics showed improved matrix effects of 49-131% in King and Ivory King salmon and were quantified with an accuracy level above 70% (**Table 3 and Table 4**). It is not yet clear which matrix components are causing these differences in matrix effects and hence accuracy.

MDLs were variable between different salmon matrices and clean-up methods. Generally, MDLs were higher in Sockeye salmon compared to the King and Ivory King salmon and clean-up methods did not show a consistent effect on the MDLs. This is likely due to different matrix effects in different salmon matrices affecting the sensitivity of the method.

## Conclusion

This work aimed to investigate a) the matrix effects from salmon on antibiotics analysis using UPLC-MS/MS, b) the effectiveness of common clean-up methods in minimizing the matrix effects, and c) potential contribution of carotenoids from salmon on the matrix effects. The QUEChERS method showed acceptable extraction recoveries but significant matrix effects which were not improved by dispersive SPE clean-up using Na_2_SO_4_/ PSA/ C18 (900/50/150) sorbents. SPE using OASIS HLB column improved matrix effects for some antibiotics but resulted in low extraction recoveries (< 30%) for most antibiotics. Carotenoids at levels found in King salmon were not major contributors to the observed matrix effects. This suggests that other co-extracts from the salmon matrix might be involved in analyte signal suppression or enhancement. Matrix effects compromised the accuracy and sensitivity of the analysis. Therefore, it is critical to characterize the nature of interfering compounds to enable better separation and accurate quantitation of antibiotics in salmon.

## Author contributions

S.E. and A.Y.T. designed the experiments. S.E. performed the experiments, analyzed the data and wrote the original draft. A.Y.T. reviewed and edited the manuscript.

## Supplementary information

The supporting information include the table presenting the MRLs for antibiotics.

### Abbreviations

AMOX: Amoxicillin
AMP: Ampicillin
ANOVA: Analysis of variance
AZ: Azithromycin
CAP: Chloramphenicol
CTC: Chlortetracycline
D4-SMZ: D4-sulfamethazine
DOX: Doxycycline
ENRO: Enrofloxacin
ENO: Enoxacin
ERYTH: Erythromycin
FF: Florfenicol
FFA: Florfenicol amine
FLU: Flumequine
HLB: Hydrophilic lipophilic Balance
LIN: Lincomycin
NOR: Norfloxacin
PEN-G: Penicillin G
PEN-V: Penicillin V
ROX: Roxithromycin
OTC: Oxytetracycline
SPE: Solid phase extraction
SDM: Sulfadimethoxine
SDZ: Sulfadiazine
SMX: Sufamethoxazole
SPE: Solid phase extraction
SSZ: Sulfasalazine
TC: Tetracycline
TAP: Thiamphenicol
TILM: Tilmicosin
TRIM: Trimethoprim
VIRG-M1: Virginiamycin M1
Virginiamycin S1: VIRG-S1.

## References

(1) Cabello, F. C. Environmental microbiology 2006, 8, 1137–1144.

(2) Done, H. Y., Halden, R. U. Journal of hazardous materials 2015, 282, 10–17.

(3) https://www.fda.gov/Food/GuidanceRegulation/GuidanceDocumentsRegulatoryInformation/Seafood/ucm150954.htm 2008.

(4) Naviner, M., Gordon, L., Giraud, E., Denis, M., Mangion, C., Le Bris, H., Ganière, J.-P. Aquaculture 2011, 315, 236–241.

(5) Kruse, H., Sørum, H. Applied and environmental microbiology 1994, 60, 4015–4021.

(6) Antibiotic / Antimicrobial Resistance (AR / AMR).

(7) Desmarchelier, A., Fan, K., Minh Tien, M., Savoy, M.-C., Tarres, A., Fuger, D., Goyon, A., Bessaire, T., Mottier, P. Food Additives & Contaminants: Part A 2018, 35, 646–660.

(8) Lopes, R. P., Reyes, R. C., Romero-González, R., Vidal, J. L. M., Frenich, A. G. Journal of Chromatography B 2012, 895, 39–47.

(9) Serra-Compte, A., Álvarez-Muñoz, D., Rodríguez-Mozaz, S., Barceló, D. Food and chemical toxicology 2017, 104, 3–13.

(10) Kruve, A., Künnapas, A., Herodes, K., Leito, I. Journal of chromatography A 2008, 1187, 58–66.

(11) Taylor, P. J. Clinical biochemistry 2005, 38, 328–334.

(12) Love, D. C., Asche, F., Conrad, Z., Young, R., Harding, J., Nussbaumer, E. M., Thorne-Lyman, A. L., Neff, R. Nutrients 2020, 12, 1810.

(13) Shamshak, G. L., Anderson, J. L., Asche, F., Garlock, T., Love, D. C. Journal of the World Aquaculture Society 2019, 50, 715–727.

(14) Turnipseed, S. B., Storey, J. M., Lohne, J. J., Andersen, W. C., Burger, R., Johnson, A. S., Madson, M. R. Journal of agricultural and food chemistry 2017, 65, 7252–7267.

(15) Storey, J. M., Clark, S. B., Johnson, A. S., Andersen, W. C., Turnipseed, S. B., Lohne, J. J., Burger, R. J., Ayres, P. R., Carr, J. R., Madson, M. R. Journal of Chromatography B 2014, 972, 38–47.

(16) Anastassiades, M., Lehotay, S. J., Štajnbaher, D., Schenck, F. J. Journal of AOAC international 2003, 86, 412–431.

(17) Rossi, R., Saluti, G., Moretti, S., Diamanti, I., Giusepponi, D., Galarini, R. Food Additives & Contaminants: Part A 2018, 35, 241–257.

(18) Islam, A. K. M. M., Hong, S.-M., Lee, H.-S., Moon, B.-C., Kim, D., Kwon, H. Journal of food science and technology 2018, 55, 3930–3938.

(19) Mandal, S., Poi, R., Ansary, I., Hazra, D. K., Bhattacharyya, S., Karmakar, R. SN Applied Sciences 2020, 2, 1–14.

(20) Authority, E. F. S. EFSA Journal 2005, 3, 291.

(21) Han, L., Matarrita, J., Sapozhnikova, Y., Lehotay, S. J. Journal of Chromatography A 2016, 1449, 17–29.

(22) Gu, H., Liu, G., Wang, J., Aubry, A.-F. o., Arnold, M. E. Analytical Chemistry 2014, 86, 8959–8966.

(23) Sapkota, A., Sapkota, A. R., Kucharski, M., Burke, J., McKenzie, S., Walker, P., Lawrence, R. Environment international 2008, 34, 1215–1226.

(24) 2016.

(25) Code of Federal Regulations, T., Volume 6, Revised as of April 1, 2018.

(26) Senta, I., Krizman-Matasic, I., Terzic, S., Ahel, M. Journal of Chromatography A 2017, 1509, 60–68.

(27) Stability of Pharmaceuticals, Personal Care Products, Steroids, and Hormones in Aqueous Samples, POTW Effluents, and Biosolids 2010.

(28) Turku, I., Sainio, T., Paatero, E. Environmental Chemistry Letters 2007, 5, 225–228.

(29) Cao, X., Pang, H., Yang, G. Journal of Soils and Sediments 2015, 15, 1635–1643.

(30) Pavlović, D. M., Ćurković, L., Grčić, I., Šimić, I., Župan, J. Environmental Science and Pollution Research 2017, 24, 10091–10106.

(31) Li, H., Yin, J., Liu, Y., Shang, J. Journal of agricultural and food chemistry 2012, 60, 1722–1727.

(32) Chamkasem, N., Lee, S., Harmon, T. Food chemistry 2016, 192, 900–906.

(33) Emerson, B., Gidden, J., Lay Jr, J. O., Durham, B. Journal of lipid research 2010, 51, 2428–2434.

