## Supplement for "Assessing the matrix effects of pigmented and non-pigmented salmon during multi-residue antibiotic analysis with liquid chromatography coupled to tandem mass spectrometry"

Table S1. Maximum residue limit (MRL; ng/g) for antibiotics according to U.S. regulations^1^.

| **Class** | **Antibiotics** | **MRL (ng/g)** | **Fish** | **Other matrices** |
| --- | --- | --- | --- | --- |
| B-Lactams | AMOX | 10 | NA | Cattle edible tissue |
|  | AMP | 10 | NA | Cattle/ swine edible tissue |
|  | PEN-G | 50* | NA | Edible tissues of cattle |
|  | PEN-V | 50* | NA | Edible tissues of cattle |
| Amphenicols | CAP | Banned | - | - |
|  | FF** | 1000 | Fish | - |
|  | TAP | NA | NA | - |
| Tetracyclines | Sum of tetracycline residues | 2000 | Finfish muscle |  |
| Quinolones | FLU | NA | NA | NA |
|  | ENO | NA | NA | NA |
|  | ENRO | 100*** | NA | Cattle liver |
|  | NOR | NA | NA | NA |
| Sulfonamides | SDZ | NA | NA | NA |
|  | SDM | 100 | Edible tissues of catfish | - |
|  | SMX | NA | NA | NA |
|  | SSZ | NA | NA | NA |
| Macrolides | ROX | NA | NA | NA |
|  | TILM | 100 | NA | Muscle of cattle |
|  | AZ | NA | NA | NA |
|  | Tylosin | 200 | NA | Muscle of cattle |
|  | VIRG | Exempt | NA | Cattle/ chicken edible tissues**** |
|  | ERYTH | 100 | NA | Cattle edible tissues |
| Lincosamides | LINǂ | 100 | NA | Swine muscle |
| Other | TRIM | NA | NA | NA |
|  | ORM | 100 | Salmonids and catfish | - |

NA: Not available

* Penicillin, MRL = 10 ng/g in turkey, 0 ng/g in chicken, milk, swine, egg, milk

**Tolerance for marker residue: FFA. Fish includes catfish muscle, freshwater-reared warmwater finfish (other than catfish) and salmonids muscle/skin

***Tolerance for desethylene ciprofloxacin (marker residue)

****Excluding cattle milk and chicken eggs; swine muscle: 100 ng/g

ǂ Exempt in chicken edible tissues

1.	Code of Federal Regulations, T., Volume 6, Revised as of April 1, 2018. <https://www.accessdata.fda.gov/scripts/cdrh/cfdocs/cfcfr/CFRSearch.cfm?CFRPart=556>.
